# Neighbor signals perceived by phytochrome B increase thermotolerance in Arabidopsis

**DOI:** 10.1101/403840

**Authors:** Denise Arico, Martina Legris, Luciana Castro, Carlos Fernando Garcia, Aldana Laino, Jorge Casal, Maria Agustina Mazzella

## Abstract

Due to the preeminence of reductionist approaches, our understanding of plant responses to combined stresses is limited. We speculated that light-quality signals of neighboring vegetation might increase susceptibility to heat shocks because shade reduces tissue temperature and hence the likeness of heat shocks. In contrast, plants of *Arabidopsis thaliana* grown under low red / far-red ratios typical of shade were less damaged by heat stress than plants grown under simulated sunlight. Shade reduces the activity of phytochrome B (phyB) and the *phyB* mutant showed high tolerance to heat stress even under simulated sunlight. The enhanced heat tolerance under low red / far-red ratios failed in a multiple mutant of PHYTOCHROME INTERACTING FACTORs. The *phyB* mutant showed reduced expression of several fatty acid desaturase (FAD) genes, proportion of fully unsaturated fatty acids and electrolyte leakage of membranes exposed to a heat shock. Activation of phyB by red light also reduced thermotolerance of dark-grown (etiolated) seedlings but not via changes in *FAD* gene expression and membrane stability. We propose that the reduced photosynthetic capacity linked to thermotolerant membranes would be less costly under shade, where the light input itself limits photosynthesis.

## 1 INTRODUCTION

Although most studies dealing with plant responses to environmental threats consider one source of stress at the time, plants can often be exposed simultaneously to multiple stresses. The impact of different stresses is not necessarily additive and in some cases one stress increases the impact of the other. For example, the damage caused by the combination of drought and salinity or drought and heat results in growth reductions that are more severe than those caused by the sum of the effects of each stress in isolation (Rizhsky, Liang & Mittler, 2002; Ahmed et al., 2013). Similarly, heat stress facilitates pathogen spread causing susceptibility to diseases (Luck et al., 2011; Nicol, Turner, Coyne, Nijs & Hockland, 2011). Conversely, in other cases one stress reduces the impact of the other. For instance, wounding can increase salt tolerance (Capiati, País& Téllez-Iñón, 2006) and ultraviolet radiation, although potentially harmful, can protect plants against herbivorous insects (Rousseaux et al., 2004; Caputo, Rutitzky & Ballaré, 2006). Therefore, plants are likely to have developed physiological and molecular mechanisms of protection against specific combinations of stresses, but these processes can remain hidden in studies involving single stress factors (Pandey, Ramegowda & Senthil-Kumar, 2015).

The shade imposed by neighboring vegetation reduces the photosynthetically active radiation available for the plants within the canopy, and can eventually compromise their survival. Plants respond to the threat associated to neighbors by inducing shade-avoidance responses (such as enhanced stem and petiole elongation), and/or acclimation responses that increase the chances of survival under limiting light (Casal, 2013; Gommers, Visser, Onge, Voesenek & Pierik, 2013). In Arabidopsis, light/shade signals are perceived mainly by phytochrome B (phyB) and cryptochrome 1 (cry1). Phytochromes are a family of five members in Arabidopsis (phyA-phyE). They have two inter-convertible forms: red light transforms the inactive Pr form into the active Pfr form, while far-red light converts Pfr back to the Pr form (Burgie & Vierstra, 2014). Therefore, the activity of phyB increases with the red / far-red ratio of the light, which is high (approx 1.1) in open places and becomes gradually depleted with the proximity of neighboring vegetation reflecting far-red light and under the canopy, which also transmits far-red more efficiently than red light (Casal, 2013; Gommers et al., 2013). There are two canonical cryptochromes in Arabidopsis (cry1-cry2), which increase their activity in response to blue light (Yu, Liu, Klejnot & Lin, 2010). phyB reduces the activity of the bHLH transcription factors PHYTOCHROME INTERACTING FACTOR 3, (PIF3), PIF4, PIF5 and PIF7 by lowering their abundance and/or DNA binding capacity (Lorrain, Allen, Duek, Whitelam & Fankhauser, 2008; Park et al., 2012). cry1 also reduces the abundance of PIF4 and PIF5 (de Wit et al., 2016; Pedmale et al., 2016). Therefore, the weaker activity of these photo-sensory receptors under vegetation shade, increases the activity of PIF3, PIF4, PIF5 and PIF7, which promote stem growth and other shade-avoidance responses (Lorrain et al., 2008; Hornitschek et al., 2012; Li et al., 2012; Leivar & Monte, 2014). The lower activities of phyB and cry1 under shade also lead to stronger nuclear accumulation of CONSTITUTIVE PHOTOMORPHOGENIC 1 (COP1), which enhances the degradation of LONG HYPOCOTYL IN FAR RED LIGHT 1 (HFR1) (Pacín, Legris & Casal, 2013; Pacín, Semmoloni, Legris, Finlayson & Casal, 2016) a negative regulator of the activity of PIFs.

High temperatures are another source of stress as they inhibit photosynthesis, damage cell membranes and cause cell death (Liu & Huang, 2000; Djanaguiraman, Boyle, Welti, Jagadish & Prasad, 2018). The expected increases in temperature over the following years predict a more severe and frequent incidence of heat stress (Hatfield & Prueger, 2015). Plants posses an inherent basal thermotolerance and also have the ability to acquire thermotolerance by the exposure to a gradual sub-lethal high temperature (heat acclimation) (Hong, Lee & Vierling, 2003). Plants cope with high temperatures by altering their physiological, morphological, biochemical and molecular status during acclimation (Bita & Gerats, 2013). For instance, plants respond to non-stressing high temperatures by increasing stem and petiole elongation and leaf hyponasty; i.e. changes that enhance leaf cooling capacity reducing the probability of stress by further temperature rises (Crawford, McLachlan, Hetherington & Franklin, 2012). Plant cell membranes are direct targets of heat stress, which increase leakage of electrolytes out of the cell (Wahid, Gelani, Ashraf & Foolad, 2007). The degree of unsaturated fatty acids in the membrane is inversely correlated with growth temperatures. A reduced proportion of polyunsaturated fatty acids in the membrane favors seedling growth at elevated temperatures (Falcone, Ogas & Somerville, 2004), but reduces seedling growth in the absence of heat stress (Routaboul, Fischer & Browse, 2000). Heat stress also promotes the immediate expression of heat-shock proteins that act to prevent and restore cell damage and preserve homeostasis (Yángüez, Castro-Sanz, Fernández-Bautista, Oliveros & Castellano, 2013; Wang et al., 2017).

The aim of this study was to investigate whether the low red / far-red ratio signals of neighboring vegetation perceived by phyB affect thermotolerance. There are several reasons that justify the proposed analysis. First, there is ecological convergence of the light and temperature cues. For instance, canopy shade reduces the irradiance and temperature levels experienced by plants (Legris, Nieto, Sellaro, Prat & Casal, 2017). Second, plant population responses to global warming can be modified by light, as in forest understory, the largest changes in thermophilization of species (the replacement of cold-adapted understory species with warm-adapted species) occurs more intensively when higher light and temperature levels coincide (De Frenne et al., 2015). Third, there is molecular convergence in the plant perception and signaling of light and temperature cues. Noteworthy, phyB functions not only as a light receptor but also as a temperature sensor, which is inactivated by far-red light and also by warm temperatures (Jung et al., 2016; Legris et al., 2016). PIF4 (Koini et al., 2009; Franklin et al., 2011; Lau et al., 2018), HFR1 (Foreman et al., 2011) and COP1 (Kim et al., 2017; Park, Lee, Ha, Kim & Park, 2017), which are described above as components of the signaling network involved in the responses to the degree of shade, play a role in thermomorphogenesis. Fourth, heat shocks modulate light signaling in etiolated Arabidopsis seedlings (Karayekov, Sellaro, Legris, Yanovsky & Casal, 2013), suggesting that the reciprocal control of thermotolerance by light signals could also occur. Fifth, a recent work has reported enhanced thermotolerance of a *phyB* mutant (Song, Liu, Hu & Wu, 2017). We found that neighbor signals perceived by phyB increase thermotolerance in Arabidopsis at least in part by adjusting membrane function. 142

## 2 MATERIALS AND METHODS

### 2.1 Plant Material

The wild-type accessions of *Arabidopsis thaliana* used in this study were Landsberg *erecta* (L*er*) and Columbia (Col). The *phyB* (*phyB-5)*(Reed, Nagpal, Poole, Furuya & Chory, 1993) and *cry1 cry2* (*hy4-2.23n, fha-1*) (Casal & Mazzella, 1998) mutants are in the L*er* background. The *phyA-211* (Reed et al., 1993), *phyB-9,*(Reed, Nagatani, Elich, Fagan & Chory, 1994), *hy5-211* (Shin, Park & Choi, 2007), *cry1-1*, *cry2-1*(Guo, Yang, Mockler & Lin, 1998), *pif1-1*, *pif3-3* (Monte et al., 2004), *pif4-101*, *pif5-3* (Lorrain et al., 2008)*, pif1 pif3*, *pif4 pif5*, *pif3 pif4*, *pif1 pif3 pif5*, *pif3 pif4 pif5*, and *pif1 pif3 pif4 pif5* (Leivar et al., 2008); *hfr-101* (Duek, Elmer, Van Oosten & Fankhauser, 2004), *cop1-4*, *cop1-6* (McNellis, 1994), *spa1-1*, *spa2-1*, *spa3-1*,*spa4-1, spa1 spa2 spa3 and spa1 spa2 spa4* (Laubinger, Fittinghoff & Hoecker, 2004) mutants are in the Col background. Seeds were surface sterilized (4 h of exposure to the fumes produced by 1.25% HCl/NaClO) and sown on 0,8% w/v agar plates (experiments with etiolated seedlings) or 0,8% w/v agar plates containing one-half-strength Murashige and Skoog basal medium pH 5.7 (MS) (experiments with light-grown seedlings). After 3 d at 4 °C in darkness, the seeds were exposed to a red-light pulse for 2 h to promote germination.

### 2.2 Light conditions

White light (WL) was provided by fluorescent lamps (Philips, 80 µmol m^-2^ s^-1^) and red light and red light (12 µmol m^-2^ s^-1^) was provided by fluorescent lamps (Philips) combined with red, orange and yellow filters (LEE #106, #105 and #101 respectively). For the WL treatments supplemented with far-red light (WL+FR), WL was given from above as described and far-red light (30 µmol m^-2^ s^-1^) was provided from below by incandescent lamps filtered with a red acetate filter and six blue acrylic filters (Paolini 2031, La Casa del Acetato, Buenos Aires, Argentina) and a water filter.

### 2.3 Temperature treatments

Plants were grown at 22 °C. For heat-shock treatments the boxes containing the seedlings were placed during 45 min in a shaker or 90 min in water bath, both at 45 °C in darkness. For acclimation to heat, the boxes containing seedlings were placed during 90 min in a shaker at 35 °C in darkness.

### 2.4 Scoring of damage

At the rosette stage, a plant was considered damaged when it showed at least one cotyledon completely bleached. At the seedling stage survival rates were assessed by recording the proportion of seedlings that generated the first pair of leaves 1 week after the heat shock.

### 2.5 Electrolyte leakage

Approximately 200 mg of the seedlings were harvested after the heat shock challenge, rinsed twice with demineralized water and subsequently floated on 10 ml of demineralized water at room temperature. Electrolyte leakage in the solution was measured 24 h later by using a conductimeter (Corning TDS-60). Data are presented relative to total conductivity obtained after boiling the samples at 100° C for 15 min (Wang et al., 2013).

### 2.6 Quantitative real time PCR (qPCR)

Total RNA was extracted using Trizol Reagent (Life Technologies Inc., USA) and treated with RQ1 RNase-free DNase I (Promega, USA). 3 µg of total RNA were reverse-transcribed in a 25 µl reaction using MMLV reverse transcriptase (Promega, USA) according to the manufacturer’s instructions, using oligo (dT) primers. cDNA were diluted 1:40 before qPCR. qPCR reactions were performed in a DNA Engine Opticon 2 System (MJ Research, USA) using the 5x HOT FIREPol EvaGreen® qPCR Mix Plus (NO ROX) kit (Solis BioDyne, Estonia). The primers for *FAD2* (AT3G12120) were 5’-CCTTCCTCCTCGTCCCTTAC-3’ and 5’-CTCTTTCGAGGGATCCAGTG-3’ and for *FAD6* (AT4G30950) were 5’-CCGTGGTATCTGCTACCGTT-3’ and 5’-TAGGAAGGCGAGAGTACCCA-3’ (Shen et al., 2010); primers for *FAD5* (AT3G15850) were 5’-AACAACTGGTGGGTAGCAGC-3’ and 5’-CTTGTAGATGCTTTCAGGCAA-3’. *ACTIN 8* (AT1G49240) was used as normalization control (Mazzella et al., 2005). The conditions for PCR were optimized with respect to primer concentrations, primer annealing temperatures and duration of steps. Cycling conditions were 95°C for 15 min followed by 38 cycles of 15 s at 94°C, 12 s at 60°C, 12 s at 72°C. PCR for each gene fragment was performed alongside standard dilution curves of cDNA pool. All gene fragments were amplified in duplicate from the same RNA preparation, and the mean value was considered.

### 2.7 Lipid Extraction and Fatty Acid Analysis

For each sample, total lipids were extracted with methanol/chloroform mix (2/1 v/v) using the procedure described by Bligh and Dyer (Bligh & Dyer, 1959). Lipid extracts were dried, weighted, suspended in 2 mL of a fresh solution of 10% KOH in ethanol and saponified for 60 min at 80 °C using stoppered glass tubes. Two ml of hexane were added and fatty acids were extracted by shaking. The upper organic phase (non-saponified) was discarded. The aqueous layer was acidified with 1.5 ml of concentrated HCl and fatty acids were extracted twice with 1.5 ml hexane. Extracts containing total free fatty acids were dried under a nitrogen stream, dissolved in 1.5 ml BF3 (10 % in methanol) and 1.5 ml benzene, and esterified by heating to 100 °C and shaking for 1 h. Fatty acid methyl ester (FAME) were extracted twice with hexane and washed with distilled water. After washing, the organic phase was evaporated under a nitrogen stream, re-dissolved in hexane, and analysed by GLC. One µl of FAME solution was injected into an Omegawax X250 (Supelco Inc., Bellefonte, PA, USA) capillary column (30 m × 0.25 mm; 0.25-mm film) in a Hewlett Packard HP-6890 (Santa Clara, CA, USA) chromatograph equipped with a flame ionization detector. The column temperature was programmed for a linear increase of 3 °C min^-1^ from 175 to 230 °C. The cromatographic peaks of FAME were identified by comparing their retention times with standards under the same conditions.

## 3 RESULTS

### 3.1 Neighbor signals increase the tolerance to a heat shock

In order to study whether neighbor signals affect thermotolerance, Arabidopsis seedlings grown for 9 d under either WL or WL+FR (high, or low red / far-red ratios, respectively) were exposed to a heat shock (45 °C for 45 min) and plant damage was recorded after 5 d of recovery at 22 °C under WL or WL+FR according to the previous growth condition (see protocol in Figure S1a). Wild-type (WT) seedlings either of the L*er* or Col background showed less frequent damage (enhanced thermotolerance) when grown under WL+FR simulating the presence of neighboring vegetation than under WL (Figure 1a,b,d,e).

**Figure 1.**
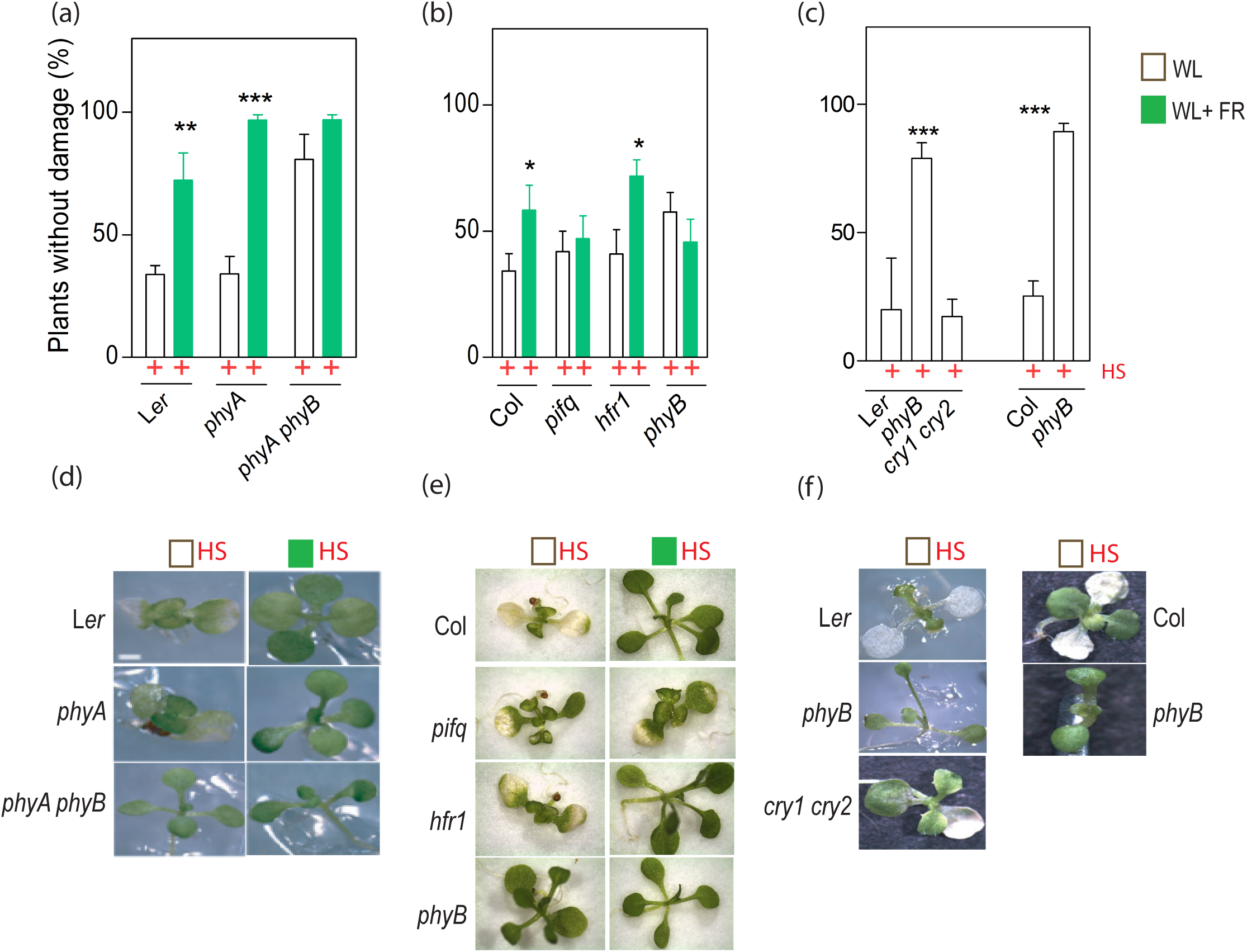
Low red / far-red ratios increase tolerance to a heat shock. Rosettes of the WT and different mutants were grown under WL or WL+FR and exposed to a heat shock (HS) during 45 min at 45 °C (protocol in Figure S1a). (a), (b), (c) Percentage of plants without damage counted 5 d after the heat shock. Data are means of at least 6 independent replicates ±SE (each replicate is average of ten plants). *, ** and *** indicate significant differences (p< 0.05, p < 0.01, p < 0.001, respectively) between WL and WL+FR (a,b) or between a mutant and the WT (c) in ANOVA followed by Bonferroni post-tests. (d), (e),) Representative photographs after the heat shock treatments. pifq: pif1 pif3 pif4 pif5 quadruple mutant.

### 3.2 phyB activity reduces the tolerance to a heat shock

In principle, the action of supplementary far-red light (i.e., WL+FR compared to WL) could be mediated by different perception and signaling steps as low red / far-red ratios reduce the proportion of active phyB but increases phyA activity (Franklin, 2003; Rausenberger et al., 2011; Trupkin, Legris, Buchovsky, Tolava Rivero & Casal, 2014) and the relative excitation of photosystems I and II (Anderson, Chow & Park, 1995). Compared to the WT, the *phyA* mutant showed no difference under WL and an enhanced response to WL+FR (Figure 1a,d). The *phyA phyB* double mutant and the *phyA* mutant showed similar thermotolerance under WL+FR and this high thermotolerance was already observed in seedlings of *phyA phyB* grown under WL (Figure 1a,d). These results indicate that in the WT, WL+FR increases thermotolerance by lowering phyB activity. Furthermore, the enhanced phyA activity caused by supplementary far-red light only slightly counteracted the promotion of thermotolerance caused by lowering phyB activity. Since the response of *phyA phyB* to WL+FR compared to WL was not significant, we obtained no evidence for a role of changes in photosystem balance (Figure 1a).

If the above conclusion is correct, the *phyB* mutation should be enough by itself to increase thermotolerance under WL (i.e. in the absence of neighbor signals). This expectation was met by the data obtained with two independent alleles (Figure 1). Under WL, the damage of the *cry1 cry2* double mutant was similar to that observed for the WT (Figure 1c,f). 274

### 3.3 Enhanced thermotolerance under low red / far-red ratios requires PIFs

Since phyB negatively regulates PIFs, the reduced activity of phyB under low red / far-red ratios increases the activity of PIFs (Lorrain et al., 2008; Li et al., 2012). Under WL,thermotolerance of the *pif1 pif3 pif4 pif5* quadruple mutant was similar to that of the WT; however, thermotolerance did not increase in *pif1 pif3 pif4 pif5* in response to WL+FR (Figure 1b,e). This suggests that in the WT, the increased thermotolerance under low red / far-red ratios is mediated by increased activity of PIFs. HFR1 is a negative regulator of the effects of PIFs on growth (Hornitschek, Lorrain, Zoete, Michielin & Fankhauser, 2009) but the *hfr1* mutation did not affect thermotolerance (Figure 1b,e).

### 3.4 phyB decreases thermostability of the plasma membranes

To test whether the increased damage observed in the WT compared to the *phyB* mutant is associated with changes in the functional integrity of plasma membranes, we evaluated electrolyte leakage in seedlings grown under WL, immediately after exposure to a heat shock of 45 min at 45 °C, compared to the seedlings that remained at 22 °C as controls (Figure S1a). Both in the L*er* and Col backgrounds, the WT, *phyB* mutants and *cry1 cry2* mutants showed similar levels of electrolyte leakage in the absence of a heat shock (22 °C) (Figure 2). After exposure to the heat shock, the WT and the *cry1 cry2* double mutant increased the leakage of electrolytes, more than the *phyB* mutant (Figure 2), indicating that phyB enhances the heat damage of the plasma membrane.

**Figure 2.**
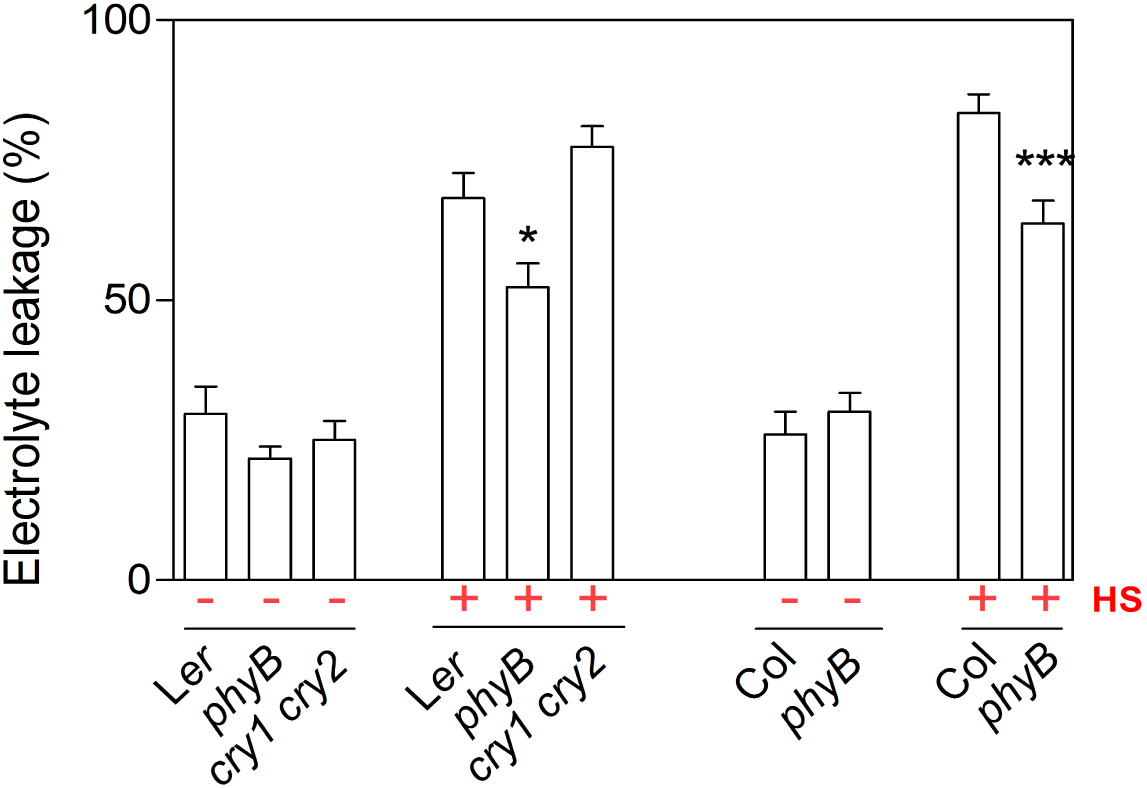
phyB increases electrolyte leakage after a heat shock. Rosettes of the WT and different mutants were grown under WL, either exposed or not exposed to a heat shock (HS) during 45 min at 45 °C and immediately harvested for the measurement of electrolyte leakage (protocol in Figure S1a). Data are the means of at least three independent replicates ±SE (each replicate is average of at least twenty plants). * and *** indicate significant differences (p < 0.05, p < 0.001, respectively) between a mutant and the WT in ANOVA followed by Bonferroni post-tests. Electrolyte leakage measurements are expressed as a percentage of the leakage achieved after boiling the plants.

### 3.5 phyB increases polyunsaturated fatty acids

Differences in membrane thermostability can result from changes in the lipid profile (Falcone et al., 2004). We therefore investigated fatty acid composition in seedlings harvested immediately prior the time when they had to be exposed to the heat shock in the above experiments (Figure S1a). Compared to the WT, the *phyB* mutant showed a significant reduction in total unsaturated fatty acids (16:3 and 18:3), a significant increase in partially unsaturated (18:1 and 18:2) but no differences in saturated (14:0 and 16:0) fatty acids (Figure 3a,b). The *cry1 cry2* double mutant showed an overall level of total unsaturated fatty acids similar to the WT (Figure 3b), an increment only in 18:2 and a slight reduction in 16:3 (Figure 3a). We are currently conducting experiments to determine unsaturation fatty acid changes in WT and *pifq mutant* shaded plants.

**Figure 3.**
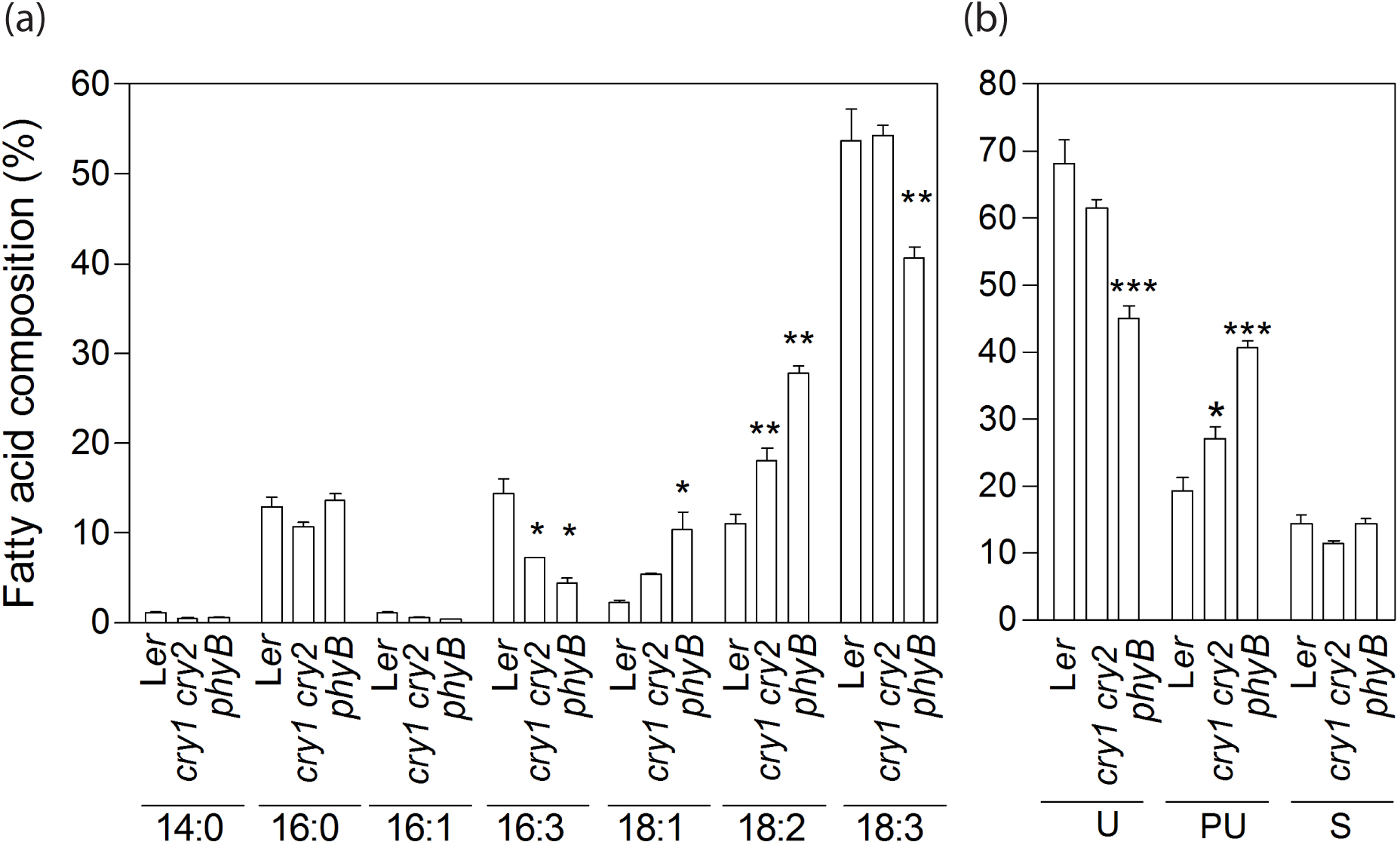
phyB increases polyunsaturated fatty acids. Rosettes of the WT and different mutants were grown under WL, and harvested at the time they would have been exposed to a heat shock (protocol in Figure S1a). (a) Total triacylglycerol fatty acid composition. (b) Saturated (S) (14:0 and 16:0), partially unsaturated (PU) (16:1, 18:2 and 18:1) and totally unsaturated (U) (16:3 and 18:3) fatty acid composition. Data are the means of at least three independent replicates ±SE (each replicate is average of ten plants). *, ** and *** indicate significant differences (p<0.05, p < 0.01, p < 0.001, respectively) between a mutant and the WT in ANOVA followed by Bonferroni post-tests.

### 3.6 phyB increases *FAD* expression

In Arabidopsis, seven fatty acid desaturase (FAD) enzymes are involved in the different steps leading to the generation of trienoic acids (Shanklin & Cahoon, 1998). Two of them, FAD2 and FAD3, localize to the endoplasmic reticulum, whereas the other five, FAD4, FAD5, FAD6, FAD7 and FAD8, localize to the chloroplast (Wallis & Browse, 2002). Given the observed changes in fatty acid composition, we evaluated whether the *phyB* mutation affects the expression of five *FAD* genes by real time PCR, in seedlings grown under WL and harvested immediately prior the time corresponding to the heat shock in above experiments (Figure S1a). The expression of *FAD2*, *FAD5*, *FAD6*, *FAD7* and *FAD8* was significantly lower in the *phyB* mutant than in the WT (Figure 4a).

At least the expression of *FAD2* and *FAD5* was reduced in WT seedlings grown under WL+FR compared to WL (Figure 4b). The response of *FAD* genes to supplementary far-red light appears to require prolonged exposures to low red / fad-red ratios because published data indicate that short-term treatments have no effects (*FAD* expression mean ± standard error. *FAD2*: WL= 12,30± 0.08, WL+FR= 12,30± 0.08; 330 *FAD5*: WL= 11,76± 0.03, WL+FR= 11,91± 0.05; *FAD6*: WL= 11,33± 0.01, WL+FR= 331 11,69± 0.02; *FAD7/8*: WL= 11,00± 0.10, WL+FR= 11,36± 0.05, data from Leivar *et al.* 332 2012).

**Figure 4.**
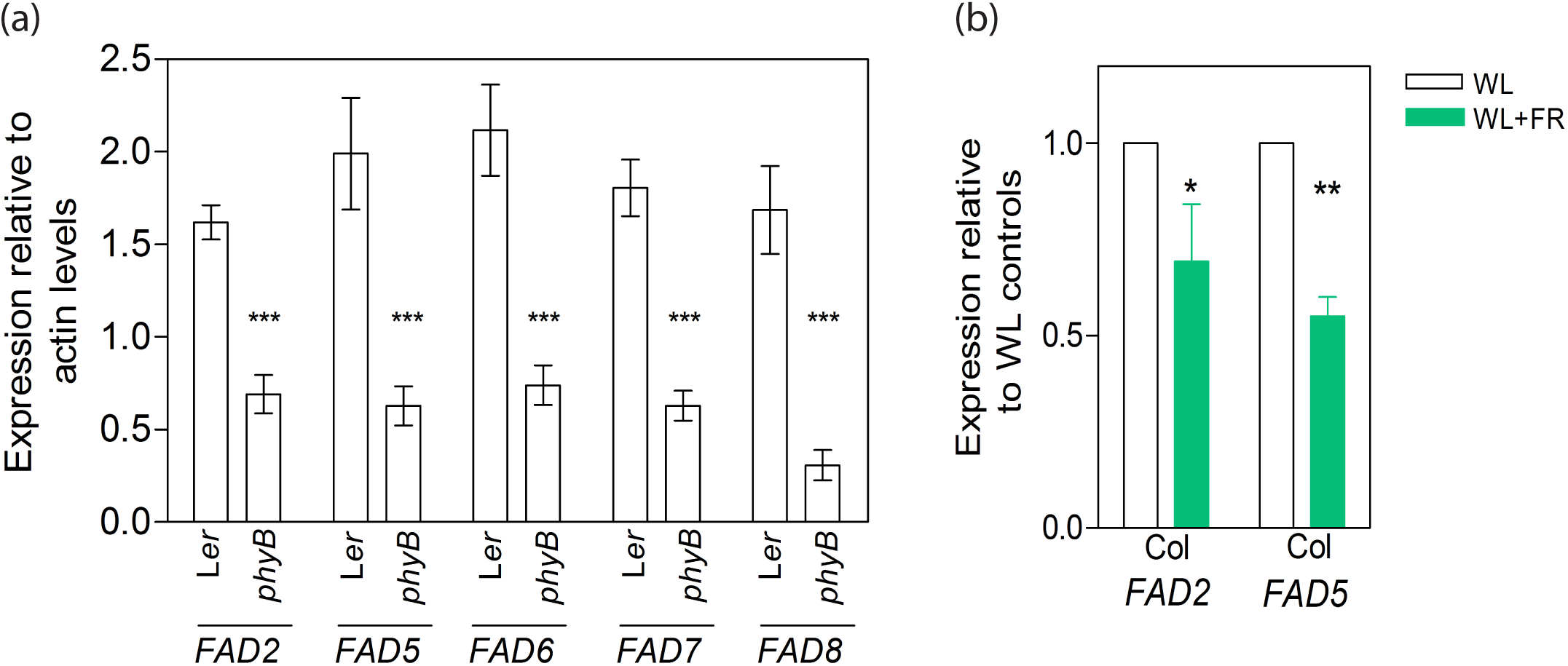
phyB increases the expression of FAD genes. Rosettes of the WT and different mutants were grown under WL or WL+FR, and harvested at the time they would have been exposed to a heat shock (HS, protocol in Figure S1a). (a) Effect of the phyB mutation under WL. Effect of WL+FR compared to WL. Data are the means of at least three independent replicates ±SE. *, ** and *** indicates significant differences (p<0.05, p<0.01 and p<0.001 respectively) in ANOVA followed by Bonferroni post-tests.

### 3.7 Light reduces thermotolerance in etiolated seedlings

To further characterize the system we analyzed the effect of phyB on thermotolerance in dark-grown etiolated seedlings. No acclimation was necessary to see a basal level of thermotolerance in light-grown seedlings (Figure 1); but in etiolated seedlings acclimation under non lethal warm temperatures was necessary. We exposed 4-day-old etiolated seedlings to a heat shock of 90 min at 45 °C followed by a 7 d recovery period before scoring survival. Prior to the heat shock the seedlings were grown for 2 d under four different conditions that resulted from the combination of a daily mild heat shock of 90 min at 35 °C to acclimate the seedlings to high temperatures and 6 h of WL (protocol in Figure S1b): the controls (no light and no acclimation treatments), the acclimated seedlings, the light-treated seedlings, and the acclimated and light-treated seedlings; although for simplicity we referred here to etiolated seedlings to those exposed to light pretreatment that were partially de-etiolated. No plant survival (0 % ±0) was observed when 4-day-old etiolated seedlings were exposed to the 45 °C heat shock without acclimation. Significant survival was observed among the seedlings that were acclimated to elevated temperatures but exposure to light during the acclimation period significantly reduced subsequent seedling survival (Figure 5a).

**Figure 5.**
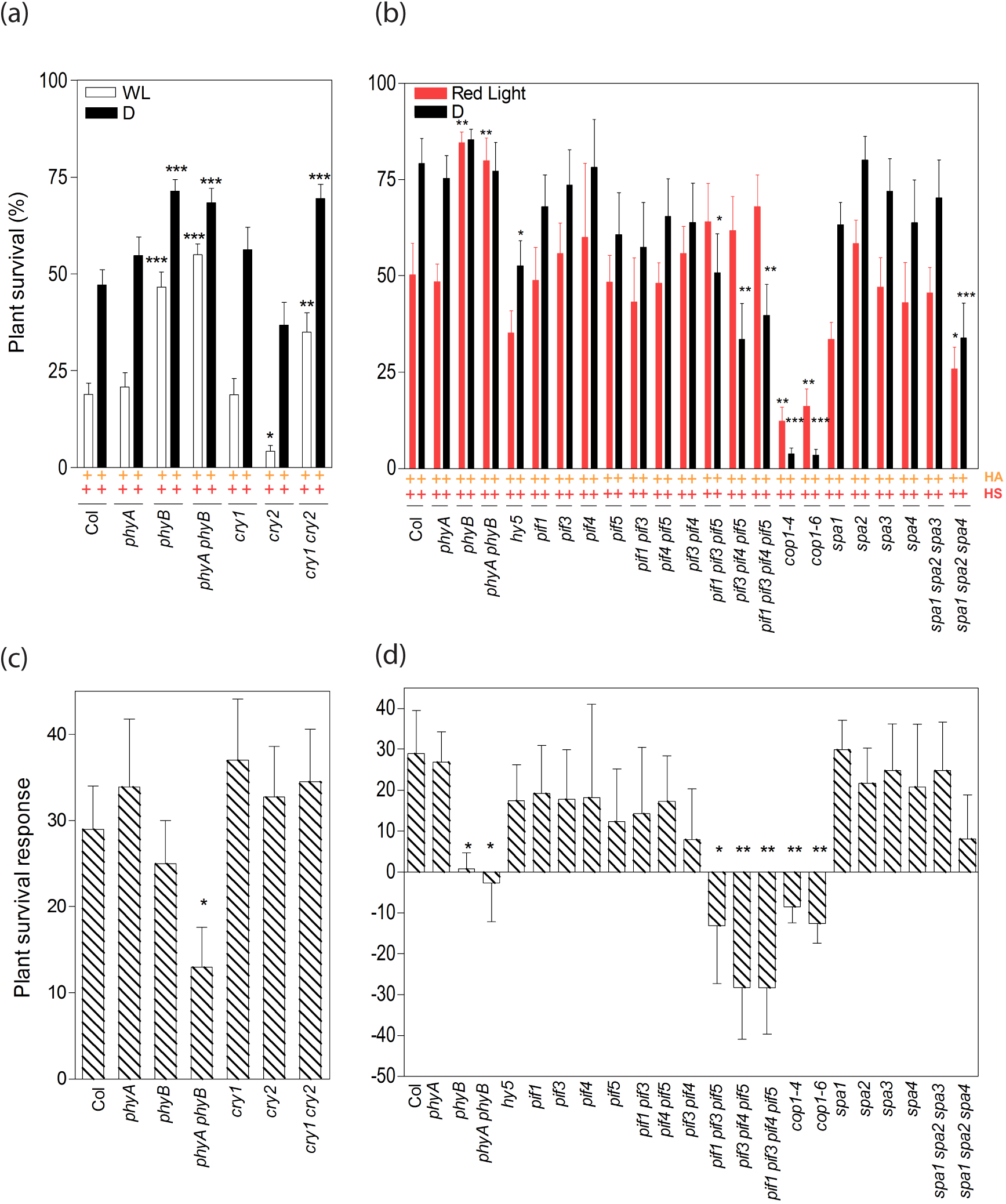
phyB reduces tolerance to a heat shock in etiolated seedlings. Etiolated seedlings were given a heat acclimation (HA) pre-treatment (35 °C) and either grown in complete darkness or exposed to light periods before a heat shock (HS) of 45 °C during 90 min followed by a recovery period (protocol in Figure S1b). Survival rates in response to WL (a), or red light (b). (c), (d) Difference in plant survival between seedlings acclimated by warm temperatures either exposed or not exposed to light. Data are the means of at least six independent replicates ±SE (each replicate is average of ten plants). *, ** and *** indicate significant differences (p<0.05, p < 0.01, p < 0.001, respectively) between a mutant and the WT in ANOVA followed by Bonferroni post-tests.

### 3.8 In etiolated seedlings, light reduction of induced thermotolerance requires phyB, PIFs, and COP1

Survival of temperature acclimated seedlings was increased in the *phyB* mutant background (*phyB* and *phyA phyB* mutants) and *cry1 cry2* double mutants with or without light treatment, while *phyA* showed the same response as the WT (note that a light treatment to induce germination was given even to the dark controls) (Figure 5a). The *cry1* simple mutants showed WT survival rates and *cry2* mutants show reduced survival, particularly when exposed to light, but the *cry1 cry2* double mutants showed increased survival (Figure 5a), indicating redundancy between cry1 and cry2 (Mockler, Guo, Yang, Duong & Lin, 1999; Mazzella & Casal, 2001). The difference in plant survival between acclimated seedlings exposed or not exposed to the light treatment is slightly reduced on *phyB* mutants and significantly reduced on *phyA phyB* mutants (Figure 5c).

Since phyB is activated by red light, we tested the same protocol but replacing the period of 6 h of WL by red light. Plant survival was reduced in the WT exposed to red light after each daily pre-treatment with warm temperature and this effect was absent in the *phyB* mutant background (Figure 5b). The plant survival response to red light was not significantly different from cero in *phyB* but retained a WT magnitude in *phyA* (Figure 5d).

Compared to the WT, in darkness, plant survival was reduced in the *hy5*, *pif1 pif3 pif5*, *pif3 pif4 pif5*, *pif1 pif3 pif4 pif5*, *cop1-4, cop1-6,* and *spa1 spa2 spa4* mutants (Figure 5b). Noteworthy, the *cop1, pif1 pif3 pif5, pif3 pif4 pif5* and *pif1 pif3 pif4 pif5* mutants showed an inverted response to light, which actually increased rather than reduced plant survival in these genotypes (Figure 5d).

### 3.9 phyB and PIFs do not affect membrane thermotolerance in etiolated seedlings

To investigate if the mechanisms that lead to a phyB-mediated reduction in heat tolerance in etiolated seedlings were similar to those described above for in fully de-etiolated seedlings, we analyzed electrolyte leakage and the expression of *FAD* genes in etiolated seedlings. The 45 °C heat shock increased leakage compared to the seedlings that did not receive this treatment, but no effects of the acclimation or light pre-treatments and of the *phyB* and *pif1 pif3 pif4 pif5* mutations were observed (Figure S2a), despite the large effects of these variants on seedling survival (Figure 5). No consistent effects of light or of the *phyB* or *pif1 pif3 pif4 pif5* mutations were observed on the expression of *FAD* genes (Figure S2b). 390

## 4 DISCUSSION

We show that *Arabidopsis* rosettes grown under low red / far red ratios, typical of places with close neighbors, are more tolerant to a heat shock than those grown under high red / far-red ratios, typical of un-shaded spots (Figure 1). The *phyB* mutant showed constitutively high thermotolerance, unaffected by the red / far-red ratio (Figure 1). Thus, the increased thermotolerance under low red / far-red ratios is mediated by a reduction in phyB activity. Although low blue light and blue / green ratios are also typical of shade and reduce cry1 and cry2 activity (Sellaro et al., 2010), the *cry1* and/or *cry2* mutations failed to enhance thermotolerance (Figure 1c,f).

The acquisition of thermotolerance under low red / far-red ratios requires PIFs because this response is lost in the *pif1 pif3 pif4 pif5* quadruple mutant (Figure 1b). Consistently, the activity of PIFs increases when that of phyB is low due either to a loss-of-function mutation or to low red / far ratios (Leivar & Quail, 2011). The *phyB* mutation reduced *FAD2, 5*, *6*, *7* and *8* transcript levels in plants grown under WL before exposure to a heat shock (Figure 4a), which could account for the lower level of polyunsaturated fatty acids in the membranes of this mutant (Figure 3). Low red / far-red ratios also decreased the expression of at least *FAD2* and *5* (Figure 4b). The activity of the *FAD7* promoter (Nishiuchi et al., 1995) and the levels of *FAD2* transcripts (Xiao et al., 2014) are enhanced by light in *Arabidopsis thaliana* and *Brassica napus*,respectively. Fatty acid desaturation is induced by red light in *Synechosystis* (Kis et al., 1998; Mironov et al., 2014). The chloroplast membrane of leaf cells contains 75-80% of unsaturated fatty acids while the membranes of non-photosynthetic tissues typically bear 60-65% of unsaturated fatty acids (McConn & Browse, 1998). Reactive oxygen species induced by heat stress promotes peroxidation of unsaturated fatty acids, damaging the membranes (Djanaguiraman, Prasad & Seppanen, 2010). Therefore, the reduced membrane damage observed in the *phyB* mutant (Figure 2) would be the result of its reduced expression of selected *FAD* genes (Figure 4) and the consequently low levels of polyunsaturated fatty acids (Figure 3). Photosynthetic reactions can increase the oxidative stress caused by heat stress (Djanaguiraman et al., 2010) but the thermotolerance phenotype of *phyB* is unlikely to result solely from its lower photosynthetic capacity (Boccalandro et al., 2009) because the *cry1 cry2* mutant also shows reduced rates of photosynthesis (Boccalandro et al., 2012) and only the *phyB* mutant exhibited enhanced thermotolerance.

In young, etiolated seedlings light-activated phyB also reduced thermotolerance (Figure 5). In both rosettes and etiolated seedlings, PIFs were required for the enhanced thermotolerance under the conditions that reduce phyB activity (low red / far-red ratios and darkness, respectively). Despite these coincidences the two scenarios showed fundamental differences. Basal thermotolerance was enough for survival of light-grown seedlings, whilst etiolated seedlings died when exposed to heat stress in the absence of warm temperature acclimation. Furthermore, in the case of etiolated seedlings no obvious effects of phyB on membrane lipid composition or electrolyte leakage in response to heat shock were observed (Figure S2). In etiolated seedlings treated as reported here, *HSFA1d* and *HSFA1e*, two master heat-shock factors (Yoshida et al., 2011; Higashi et al., 2013) showed increased expression in response to warm temperature acclimation in the dark and this effect was significantly reduced by light (Karayekov et al., 2013). Real time PCR of these heat shock factors on our samples did not allow us to arrive to a conclusive result, possibly because of the low expression levels of these genes. However, taking into account the results of Karayekov et al, the plausible working hypothesis that in etiolated seedlings phyB activity reduces the tolerance to a heat shock by lowering the expression of *HSFA1d* and *HSFA1e*, induced by warm temperature acclimation, would require further deep studies.

In tomato, low phyB activity due either to far-red light or to the loss-of function *phyB* mutation increases cold tolerance by increasing *CBF1* transcript levels (Wang et al., 2016). In *Oryza sativa* the *phyB* mutant shows reduced membrane lipid peroxidation and increased membrane integrity in response to cold stress (He et al., 2016). Taken together with the enhanced tolerance to heat stress reported here and elsewhere (Song et al., 2017), these observations indicate that the *phyB* mutant is less susceptible to both temperature extremes. Several abiotic stress gene markers, including heat-stress genes, are expressed at higher levels in the *phyB* mutant, which would be more resistant to abiotic stress (Yang, Seaton, Krahmer & Halliday, 2016).

Since radiation load is one of the main controls of the temperature of plant tissues (Legris et al., 2017), heat stress would be more likely to affect plants fully exposed to sunlight than plants shaded by neighboring vegetation. Furthermore, heat stress can be more severe if combined with high light, due to enhanced photo-oxidative stress (Foyer, Descourvières, Kunert & Descourvieres, 1994). Therefore, at first glance the observation that tolerance to heat stress is higher in plants grown under low red / far-red ratios typical of shade is counterintuitive. However, a more complete picture is obtained if not only the probabilities of heat stress but also the costs of thermotolerance are taken into account. Photosynthesis is favored by membranes relatively reach in unsaturated fatty acid (McConn & Browse, 1998), which are more susceptible to heat stress (Routaboul, Skidmore, Wallis & Browse, 2012). Therefore, for a plant grown under high red / far-red ratios, low thermotolerance might help to optimize the use of the light resource unless warm temperatures anticipate the likely occurrence of a heat shock and induce acclimation. Under the low red / far-red ratios of shade, photosynthesis is limited by light availability and therefore, reducing the level of unsaturated fatty acids would not come at a significant cost because photosynthesis is already reduced. In tomato, low red / far-red ratios reduce the photosynthetic capacity of the stem and the rate of respiration of this organ, in a response that saves energy when light capture is compromised by shade (Cagnola, Ploschuk, Benech-Arnold, Finlayson & Casal, 2012). Along the same line, the abundance of proteins involved in chloroplast biogenesis is reduced in multiple photoreceptor mutants (Fox, Barberini, Ploschuk, Muschietti & Mazzella, 2015). Furthermore, phytochrome mutants are less affected by growth-restricting abiotic stresses but this comes at the cost of reduced growth in the absence of stress (Yang et al., 2016). The emerging picture is that phyB activity could act as a switch between a growth-promoting status to take advantage of the resources in open places and a more conservative, stress-tolerant mode when the resources for growth become limited by competition with neighbors.

## 5 ACKNOWLEDGMENTS

This work was supported by *Agencia Nacional de Promoción Científica y Tecnológica*, Argentina (PICT 2014-545 to MAM, and PICT-2016-1459 to JJC); and CONICET Argentina (PIP 2013-2015 num 455 to MAM). We thank Dr. Romina Fox for technical support with real time PCRs.

